# Wavelength-dependent effects of artificial light at night on phytoplankton growth and community structure

**DOI:** 10.1101/2021.02.08.430211

**Authors:** Christina Diamantopoulou, Eleni Christoforou, Davide M. Dominoni, Eirini Kaiserli, Jakub Czyzewski, Nosrat Mirzai, Sofie Spatharis

## Abstract

Artificial light at night (ALAN) is increasingly recognised as a disruptive form of environmental pollution, impacting many physiological and behavioural processes that may scale up to population and community-level effects. Mounting evidence from animal studies show that the severity and type of the impact depends on the wavelength and intensity of ALAN. This knowledge has been instrumental for informing policy-making and planning for wildlife-friendly illumination. However, most of this evidence comes from terrestrial habitats, while research testing alternative wavelength illumination in marine environments is lagging behind. In this study we investigated the effect of such alternative ALAN colours on marine primary producers. Specifically, we tested the effect of green, red, and natural white LED illumination at night, compared to a dark control, on the growth of a green microalgae as well as the biomass, diversity and composition of a phytoplankton assemblage. Our findings show that green ALAN boosted chlorophyll production at the exponential growth stage, resulting in higher biomass production in the green algae *Tetraselmis suesica.* All ALAN wavelengths affected the biomass and diversity of the assemblage with the red and green ALAN having the stronger effects, leading to higher overall abundance and selective dominance of specific diatom species compared to white ALAN and the dark control.

**Synthesis:** Our work indicates that the wavelength of artificial light sources in marine areas should be carefully considered in management and conservation plans. In particular, green and red light should be used with caution in coastal areas, where there might be a need to strike a balance between the strong effects of green and red light on marine primary producers with the benefit they bring to other organisms.

## Introduction

During the last century the use and misuse of artificial light at night (ALAN) has increased considerably. Recent analyses have suggested that ALAN, which is strongly associated with the increasing worldwide urbanisation (Seto, Güneralp, & Hutyra, 2012), is still currently spreading spatially at a rate between 2 and 6% per year (Hölker et al., 2010; Kyba et al., 2017), with a parallel increase in irradiance at 2.2 % per year (Kyba et al., 2017). The surge of ALAN has altered natural lightscapes, which in turn may have dramatic effects on wild species and ecosystems. Indeed, the impact of ALAN on wildlife and ecosystems has received a lot of attention in the last two decades (D. Dominoni, Quetting, & Partecke, 2013; Gaston, Davies, Bennie, & Hopkins, 2012; Longcore & Rich, 2004; Navara & Nelson, 2007). In vertebrates, ALAN has been linked to several behavioural and physiological effects, such as disruption of circadian rhythms (Gaston, Davies, Nedelec, & Holt, 2017), altered reproductive timing (D. Dominoni et al., 2013; Robert, Lesku, Partecke, & Chambers, 2015), poor sleep (Aulsebrook et al., 2020; Raap, Pinxten, & Eens, 2015), reduced immune function (Kernbach et al., 2019) and altered metabolism (Pulgar et al., 2019). Insects are also heavily affected (Knop et al., 2017; Owens et al., 2020; van Grunsven et al., 2020), particularly because of the strong phototaxis found in many species (Altermatt & Ebert, 2016; van Langevelde et al., 2018).

Despite the surge of interest in the ecological effects of ALAN, most of the evidence collected so far comes from terrestrial habitats, while studies on marine species and communities are currently limited (Davies, Duffy, Bennie, & Gaston, 2014; Fobert, Da Silva, & Swearer, 2019; Grubisic, 2018; Witherington & Bjorndal, 1991). In particular, studies on primary marine producers, such as phytoplankton, are currently lacking. Due to the continuous expansion of coastal urbanisation (Henderson et al., 2020; Yi, Qian, Kobuliev, Han, & Li, 2020), artificial light at night is a source of pollution that is increasingly relevant for coastal ecosystems (Davies et al., 2014). Coastal ecosystems globally are also increasingly affected by eutrophication and harmful algal blooms due to nutrient-rich inflows from either agricultural or urban sources (Justic, Rabalais, Turner, & Diaz, 2009; Spatharis, Danielidis, & Tsirtsis, 2007). Given the ecological importance of light for photoautotrophic phytoplankton species, the potential severity of ripple effects from phytoplankton to higher trophic levels, and the existing vulnerability of coastal systems to eutrophication, it is imperative to determine the type and magnitude of the response of marine primary producers to ALAN.

In primary producers, light is a strong modulator of photosynthesis and associated processes driving growth and cell fitness (Edwards, Thomas, Klausmeier, & Litchman, 2015). Light absorption, by chlorophyll, peaks at approximately 430 nm, although a second, lower peak is also present at longer wavelengths of 670 nm (Grubisic et al., 2017; Lohrenz, Weidemann, & Tuel, 2003; Luimstra, Verspagen, Xu, Schuurmans, & Huisman, 2020). Moreover, light serves as an informational signal, regulating the synchronization of diverse intracellular processes ranging from phototactic, photoprotective and physiological responses essential for growth and development (Duanmu, 2017; Serrano-bueno, Romero-campero, Lucas-reina, Romero, & Valverde, 2017; Wobbe, Bassi, & Kruse, 2016). For example, green algae have an “eyespot” whereby, through a rhodopsin mediated signalling pathway, are able to direct their movement (Hartmann Harz & Hegemann, 1992; Sineshchekov & Govorunova, 1999). Therefore, the disruption of light cycles by ALAN has the potential to impact the physiology, and consequently the assemblage structure, of these organisms via multiple pathways that are responsive to different light wavelengths. This is a topical question because the spectral composition of ALAN is also changing along its surge in intensity, since many countries are replacing traditional lighting sources with the cost-efficient, energy-saving light-emitting diode (LED) technology (Gaston et al., 2012). LEDs are very flexible light sources whose colour can be easily modified. Indeed, new light installations use LEDs of different colours. While cool white LEDs are the most widespread, natural-warm white, green and red LEDs are also in use (Davies et al., 2014; Gaston et al., 2012).

Recent findings have demonstrated that ALAN from warm white High Pressure Sodium (HPS) lamps (rich in yellow/orange/red wavelengths) can affect multiple signalling events and metabolic pathways essential for photosynthesis in freshwater cyanobacteria (Poulin et al., 2014). ALAN from cool white LEDs (richer in blue/green wavelengths) can increase the photosynthetic biomass of microphytobenthos (Maggi & Benedetti-Cecchi, 2018) and its temporal variability (Maggi, Bertocci, & Benedetti-Cecchi, 2020), alter periphyton composition (Grubisic, 2018) and modify community structure of freshwater benthic microorganisms (Hölker et al., 2015). However, the aforementioned studies experimented with a single ALAN wavelength, while it is increasingly recognised that different wavelengths can cause profoundly different responses in wild organisms (Brüning, Hölker, Franke, Kleiner, & Kloas, 2016; D. M. Dominoni et al., 2020; Donners et al., 2018; Gaston, Bennie, Davies, & Hopkins, 2013; Longcore et al., 2015; Ulgezen et al., 2019). This can potentially lead to competing conservation goals (Davies, Bennie, Inger, de Ibarra, & Gaston, 2013). In coastal areas, red LED light has been recommended as a source of illumination because it doesn’t interfere with sea turtle nesting and hatching (Miller and Bretschneider 2006), as well as with coral biology (Ayalon, de Barros Marangoni, Benichou, Avisar, & Levy, 2019), whereas green light was suggested to minimise the impact of ALAN on seabird navigation (Poot et al. 2008). However, studies on the effects of ALAN wavelengths on coastal ecosystems are scarce. To understand and therefore inform the ecological management of coastal areas, it is essential to establish the effects of different wavelengths of ALAN on important aspects of microalgae assemblages, ranging from single species responses to community level properties such as diversity and species composition.

In the present study we investigate experimentally the response of marine phytoplankton to three ALAN wavelengths [natural white 4500K (470 nm), green (525 nm) and red (624 nm)] compared to dark nights. Our first objective is to determine whether different wavelengths of ALAN can have different impact on the growth of a single phytoplankton species. Furthermore, we aim to assess whether different wavelengths can have different impacts on the diversity and species composition of a phytoplankton assemblage. We hypothesize that white ALAN might stimulate growth compared to the dark, as it partly overlaps in wavelength with the first absorption peak of chlorophyll-a, an abundant pigment in all microalgae, at approximately 465 nm (Luimstra et al., 2020). Conversely, we predict that the green and red ALAN should have a weaker, if any, effect as its spectral properties have a minimal overlap with the light absorption range of chlorophyll-a (Figure S1). Finally, we predict that the effect on single species growth could cascade to the community level, as phytoplankton species’ competitive ability has been shown to shift with water colour (Luimstra et al., 2020).

## Materials and methods

### Experimental set up and light sources

To test for the effect of ALAN wavelengths on single species growth and assemblage biomass and diversity we run two concurrent experiments from 23/11/2021 and for a period of 18 days: Experiment 1 tested the effects of ALAN on the green microalgae *Tetraselmis suesica* and experiment 2 on a diatom-dominated natural coastal assemblage. The experimental design comprised of four treatments: control (12:12 Light-Dark), and three different ALAN wavelengths, green, red and natural white (12:12 Light-ALAN). Each of the two experiments comprised of five replicated cultures within each of the four treatments for a total of 40 experimental units (Erlenmeyer flasks of 200ml each).

Since algae use light as a source of both information and energy (Bennie, Davies, Cruse, & Gaston, 2016; Falcón et al., 2020), light treatments were standardised to irradiance levels rather than illuminance levels. Daytime light irradiance was 6.5 Watt m^−2^ and was provided by a 10W flood light (Prolite, Ritelite Systems Ltd, UK) equipped with two arrays of high-power LEDs (6,000K). Each ALAN source consisted of a strip of 3 LED diodes. The green ALAN wavelength was 525 nm (MULTICOMP), the red was 624 nm (MULTICOMP) and the broad-spectrum white LED light contained a higher peak at 470 nm and lower peaks between 550-600 nm (BROADCOM) (for full spectral characteristics of LED lights see Fig. S1). The emission spectra were measured by a spectrometer (AvaSpec-2048L, Avantes, Apeldoorn, The Netherlands). Night-time light irradiance was measured with a LI-200R pyranometer (LI-COR, USA) and was standardised at 0.023 Watt m^−2^ for all three ALAN treatments. This irradiance level is within the range of values reported in previous studies on ALAN (Grubisic, van Grunsven, Manfrin, Monaghan, & Hölker, 2018; Hölker et al., 2015; Levy et al., 2020; Tamir, Lerner, Haspel, Dubinsky, & Iluz, 2017).

The distance between the surface of the water in the flasks and the LED lights was approximately 40 cm. Each light treatment was applied inside a light-proof box (55×62×62cm), where the experimental replicates were introduced (see below for details on how these were produced in each experiment). The replicates were partially submerged (by 1/3 of the flask height) into water baths (44×41×22cm). Light irradiance was measured at six different locations within each water bath, and was not statistically different between the ALAN treatments both during the day (linear mixed model, χ = 3.9, p = 0.41) and during the night (linear mixed model, χ = 4.1, p = 0.13). Temperature in the water baths was maintained at a temperature of 14-15°C chosen to reflect the mean annual sea surface temperature of mid-latitude seas. To minimise box effects, water in the baths had identical temperature as it was fed from a central tank were temperature was regulated. Cultures were mixed once a day when their position inside each treatment box was also randomised.

### Experimental procedure for experiment 1: single species response

The green microalgae *T. suesica* was selected for the single species response experiment because of its use as a model species in studies using continuous illumination with monochromatic LEDs (Abiusi et al., 2014; Aidar et al., 1994; Schulze et al., 2016), because of the industrial potential of the species as a high-lipid content strain (Montero et al. 2011), and thus its importance as fish and shellfish aquaculture feed (Muller-Feuga, 2013).

Inoculum from our *T. suesica* culture (sourced by CCAP 66/4) was grown in F/2 medium (Guillard 1975) made by ultrapure artificial seawater at 35ppm salinity (V= 200mL). All cultures were initiated at a concentration of 5,000 cells/mL. Every second day, 5ml samples were taken from each replicated culture, two hours after the onset of day light in the morning, to calculate cell numbers and growth rate. Cells were counted using Fast-Read® 102 counting chambers under a light microscope. *T. suesica* growth showed a lag phase of 8 days due to the acclimation of cells from 20°C to 15°C and thereafter growth entered the exponential phase. Maximum growth rate for each replicate culture was determined based on the formula μ = ln(N_2_/N_1_)/(t_2_ - t_1_), where μ is the specific growth rate, and N1 and N2 are the cell number at time 1 (day 8) and time 2 (day 18), respectively. At day 18 of the experiment, 50ml samples were also taken to determine chlorophyll-a concentration according to Parsons et al. (1984).

### Experimental procedure for experiment 2: diatom assemblage response

We used a natural diatom-dominated marine sample in our assemblage response experiment. Specifically we assorted, at equal volume (200ml) across the 20 experimental units, an inoculum of unfiltered marine surface water collected at 50cm depth from the shore of Largs, Scotland (55.794659, −4867615). The initial inoculum had a chlorophyll-a concentration of 0.7 μg/L and salinity 30 psu which is lower than the salinity of the open sea as the area receives freshwater inflows.

The culture medium consisted of the collected marine sample and added nutrients commensurate with F/2 medium concentration (Guillard, 1975). Species identities, cell counts and chlorophyll concentration were determined on day 12 of the experiment when cultures just entered the stationary phase as determined by the cell counts of selected replicates. Specifically, a 5 ml sample was collected for species identification and was preserved with Lugol’s iodine solution. Samples were subsequently filtered through a SartoriusTM Cellulose Nitrate Membrane Filters (0.45μm pore size, 25mm diameter) and dried in an incubator at 40°C for an hour. The filter was made transparent by the addition of a drop of immersion oil and was observed under a light microscope (40x/0.65) where 15 randomly-selected fields of view were used to identify and enumerate the different species. The volume of sample examined was equal across all samples thus species’ cell counts as well as total assemblage cell counts are directly comparable across replicates and reported as counts. Chlorophyll concentration was determined from 50 ml samples as in the case of the *T. suesica* experiment.

### Data analysis

For the *T. suesica* single species response, we used three Gaussian linear models to determine the effect of ALAN treatment (4 levels: green, red and white ALAN and the dark control) on each of three response variables: the growth rate, and the cell number and chlorophyll-a measured on the final day of the experiment (day 18).

For the diatom assemblage response, we used four linear models to test the effect of ALAN treatment on each of four response variables: assemblage total cell count, chlorophyll-a, Menhinick richness and Pielou’s evenness (Pielou, 1966). The Menhinick species richness index was used to enable standardisation of species richness across samples based on the total cell abundance. This index was shown to be sensitive in expressing changes in phytoplankton diversity by previous studies comparing multiple diversity indices using phytoplankton species abundance data (Spatharis & Tsirtsis, 2010). Assemblage total cell count, chlorophyll-a and Menhinick richness were modelled with Gaussian models. Evenness was modelled with a beta distribution model as its values were confined between 0 and 1 and the effect of treatment was tested by model selection using likelihood ratio test.

To test the effect of ALAN treatment on assemblage composition, we performed analysis of similarity between all pairwise combinations of the 20 replicates using the Bray-Curtis similarity index (Arhonditsis, Karydis, & Tsirtsis, 2003) on non-transformed species-abundance data. We visualised these similarities using cluster analysis in order to check the grouping of samples based on the different treatments. We fitted additional linear models to test for the effect of treatment on the abundance of specific diatom species. Finally, the percentage changes we report in the first paragraph of the discussion were calculated according the formula: [(treatment 1-treatment 2)/treatment 1]*100.

All models were fitted based on the least squares approach apart from the evenness model that was fitted with log likelihood. We also conducted post-hoc pair-wise t-tests to assess differences between the four treatment levels. All statistical analyses was carried out in R v.3.5.0 (RStudio Team, 2016). The packages ggplot2 v.3.3.0 (Wickham et al., 2020), ggpubr v.0.2.5 (Kassambara, 2020), ggdendro v0.1-20 (de Vries & Ripley, 2016) and dendextend v.1.13.4 (Galili, 2015) were employed for plot generation and data visualisation. The package emmeans was used for pairwise comparisons between treatment levels (Lenth, 2018). For data manipulation, reshape2 v.1.4.3 (Wickham, 2007), plyr v.1.8.6 (Wickham, 2011). For modelling with the beta distribution we used the package glmmTMB (Brooks et al., 2017). We used the *vegan* v.2.5-6 R (Oksanen et al., 2019) and cluster v.2.1.0 packages (Rousseeuw et al., 2019) to perform the pairwise similarity of species-abundance data and related cluster analysis.

## Results

### Green and red ALAN promote growth of the green microalgae Tetraselmis suesica

The ALAN treatments had a statistically significant effect on the growth rate of *T. suesica* cells (F_3,16_ = 6.64, p =0.004) and this was shown to be wavelength specific. Specifically, a significantly higher growth rate was observed under the green ALAN treatment compared to white ALAN and the dark treatment. Furthermore, the red ALAN was also higher than the white ALAN treatment (Fig. 1A) (see Supplementary Table 1 for results of post-hoc tests).

**Figure 1.**
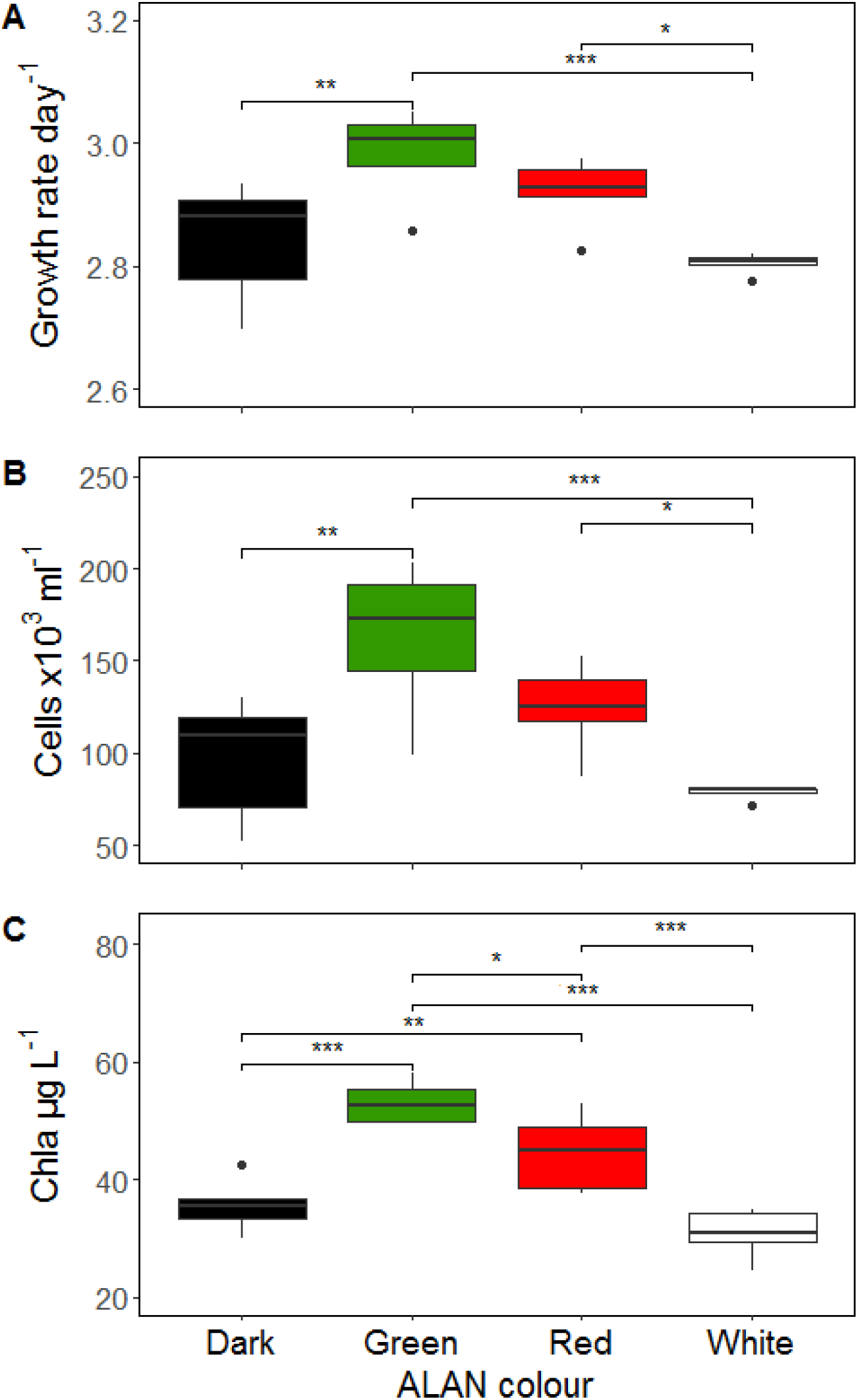
ALAN affects growth rate, cell and chlorophyll-a concentration of *Tetraselmis suesica* in a colour-dependent manner. Effect of ALAN treatments (Dark, Green, Red and White) on the growth rate calculated during the exponential growth phase (days 8-18) (panel A), on the cell concentration at day 18 (panel B) and chlorophyll-a concentration at day 18 (panel C). Pairwise comparisons show differences between treatments (not shown: p > 0.05, *: p <= 0.05, **: p <= 0.01, ***: p <= 0.001).

The *T. suesica* cell concentration was significantly affected by the ALAN treatment (F_3,16_ = 7.691, p<0.002). Specifically, on day 18 of the experiment, the cell number was significantly higher in response to the green ALAN treatment compared to the white ALAN and dark treatments, and was also higher in the red ALAN compared to white. No difference was observed between the white and dark treatments (Fig. 1B, see also Supplementary Table 1 for results of post-hoc tests). These results are comparable to those obtained in a pilot experiment where LED colours were allocated to different experimental boxes and light intensity was standardised at 20 lux (for details of this pilot experiment and relative results, see Fig. S2).

The chlorophyll-a concentration of *Tetraselmis* cultures on day 18 was also significantly affected by the ALAN wavelength (F_3,16_ = 20.584, p<0.001). Specifically, chlorophyll-a content was significantly higher in the red and greed ALAN treatments compared to the dark and white treatments, whereas no difference was observed between the white and dark treatments (Fig. 1C, see also Supplementary Table 1 for results of post-hoc tests).

### ALAN affects assemblage biomass and assemblage evenness

Analysis of the initial inoculum upon collection from the sea showed that the assemblage was comprised of 14 species of which 11 species were Diatomophyceae, two were Dinophyceae and one was Dictyochophyceae. Most dominant species were *Skeletonema* sp. (28% dominance), *Cyclotella* sp.1 (19% dominance), *Cyclotella* sp.2 (17% dominance), *Ceratium lineatum* (15% dominance) and *Navicula* sp.1 (6% dominance) whereas all other species were subdominant with relative abundance <2%. On day 12 of the experiment, overall biomass had considerably increased and stabilised across treatments and assemblages. Experimental units on day 12 comprised of 4-7 species of diatoms (13 species overall across all treatments). The planktic colonial diatoms *Skeletonema* sp., *Thalassiosira nordenskioeldii,* and to a lesser extend *T. eccentrica* were more dominant across treatments. However, the absolute and relative abundance of *Skeletonema* sp., *T. nordenskioeldii* presented differences between treatments as discussed below.

ALAN affected assemblage cell counts and diversity independent of colour whereas chlorophyll-a concentration was affected in a wavelength specific manner. Specifically, the total diatom assemblage cell count (i.e., cells summed across all species in the assemblage) was significantly affected by every ALAN treatment tested (F_3,16_=9.589, p< 0.001). Cell count was statistically higher under all ALAN wavelength conditions compared to the dark but no differences were observed between the ALAN wavelengths (Fig. 2A, see also Supplementary Table 2 for results of post-hoc tests). The chlorophyll-a concentration of the diatom assemblage was also significantly affected by the variable treatment (F_3,16_=12.393, p< 0.001), with all ALAN wavelengths having a higher concentration compared to the dark control whereas further differences were observed between the red wavelength from the green and white (Fig. 2B, see also Supplementary Table 2 for results of post-hoc tests). A significant effect of treatment was also observed on the assemblage evenness (LRT if dropped, DF=3, p<0.001), whereby the assemblages under all ALAN wavelengths had significantly higher evenness (more evenly distributed species’ populations) compared to the dark control (Fig. 2C, see also Supplementary Table 2 for results of post-hoc tests). A significant effect of ALAN was observed on the Menhinick richness (F_3,16_=3.260, p=0.049), with the dark treatment showing significantly higher richness compared to all ALAN wavelengths tested (Fig. 2D, see also Supplementary Table 2 for results of post-hoc tests).

**Figure 2.**
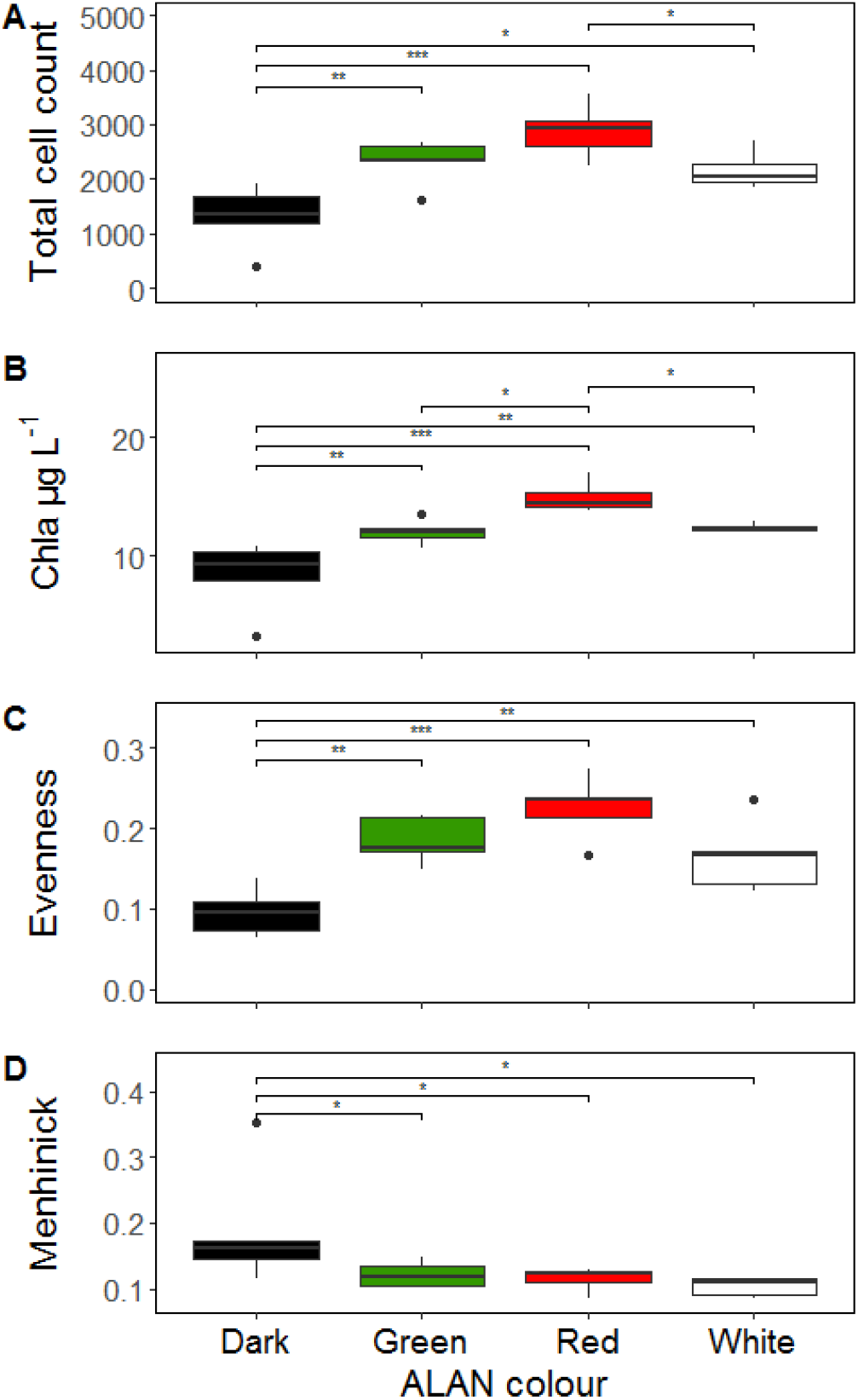
ALAN affects assemblage biomass and diversity. Effect of treatment (dark and green, red and white) on the total cell counts (panel A), chlorophyll-a (panel B), evenness (panel C) and Menhinick richness measured on day 12 of the diatom assemblage experiment. Pairwise comparisons show differences between treatments (not shown: p > 0.05, *: p <= 0.05, **: p <= 0.01, ***: p <= 0.001).

### ALAN affects species’ relative abundance

ALAN did not lead to a shift in the species’ identities comprising the assemblages. However, ALAN affected the absolute and relative abundance of species (measured as standardised cell counts) within treatments in a wavelength specific manner. In particular, assemblages growing under the green and red ALAN were 80% similar and were 44% dissimilar from the assemblages under the dark control and natural white ALAN conditions (Fig. 3A). This was due to a significant increase in the species *Skeletonema* sp. and *T. nordenskioeldii* relative to the subdominant species in the assemblage (i.e. all species excluding *Skeletonema* sp., *T. nordenskioeldii* and *T. eccentrica).* Specifically, *Skeletonema* sp. had a significantly higher abundance in all ALAN colours compared to the control (F_3,16_=7.708, p=0.002) (Fig. 3B). *T. nordenskioeldii* was significantly higher in response to the red ALAN treatment compared to the natural white and the dark control (F_3,16_=8.8574, p=0.001) (Fig. 3C). No differences between the treatments were observed in the abundance of the subdominant species (F_3,16_=0.833, p=0.495) (Fig. 3D).

**Figure 3.**
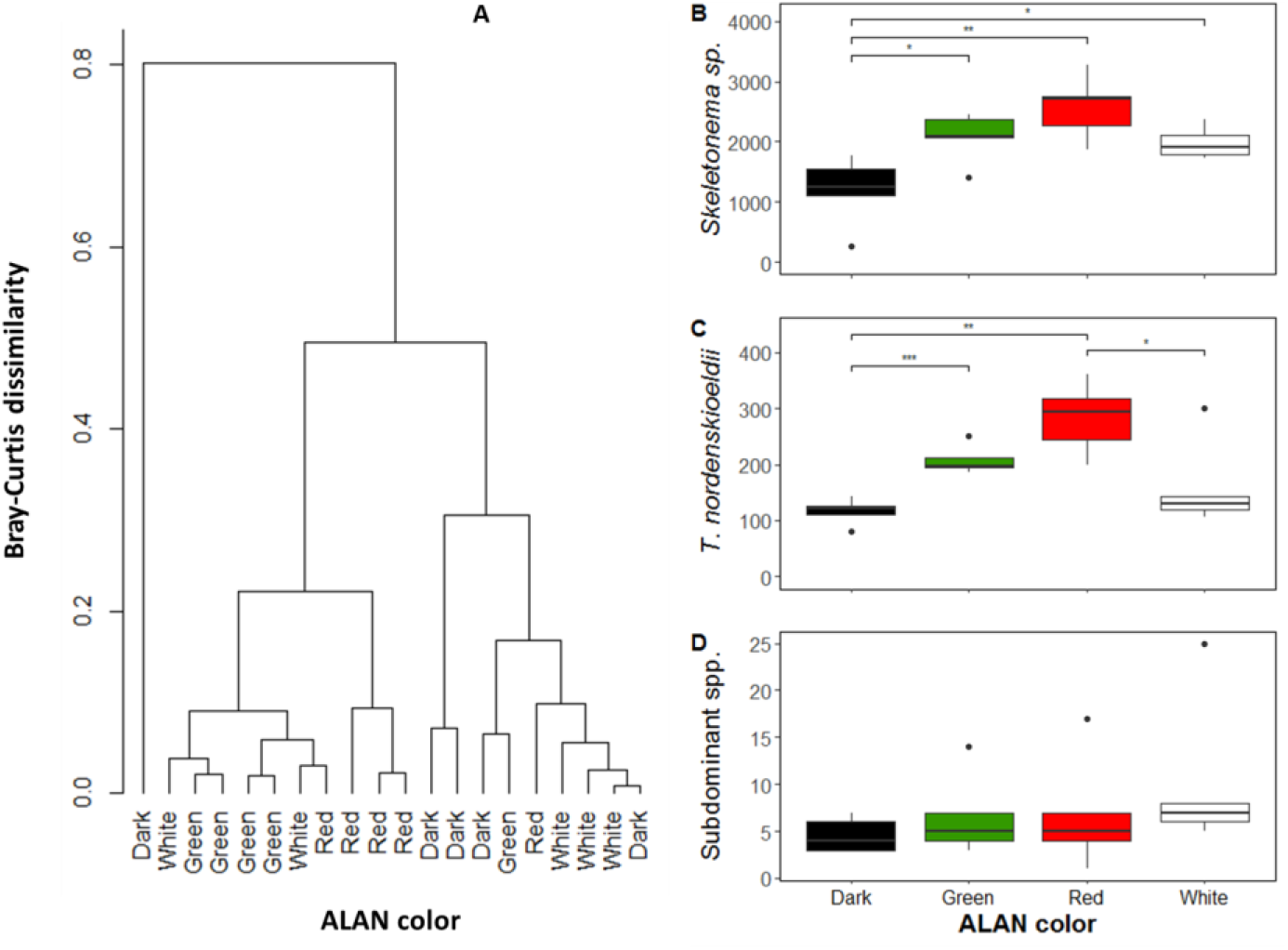
Red and green ALAN lead to similar responses in assemblage composition. Cluster showing the pairwise similarities between the replicate assemblages based on the Bray-Curtis similarity index calculated on non-transformed species-abundance data (panel A). Pairwise comparisons of abundances of *Skeletonema* sp. (panel B)*, Thalasiosira nordenskioeldii* (panel C), the sum of all subdominant species in the assemblage (panel D) (not shown: p > 0.05, *: p <= 0.05, **: p <= 0.01, ***: p <= 0.001).

## Discussion

In this study, we tested the effect of different ALAN wavelengths on phytoplankton growth, assemblage diversity and species composition. We predicted that the effect on single species and assemblage level would be more pronounced under white ALAN, compared to dark and red and green ALAN, as it partly overlaps in wavelength with the first absorption peak of chlorophyll-a, an abundant pigment in all microalgae (Luimstra et al., 2020). Contrary to our expectations, our findings suggest that red and green ALAN have more pervasive impact on phytoplankton growth and assemblage structure compared to the natural white ALAN. More specifically, our experiments showed that exposure of the green microalgae *Tetraselmis suesica* to green ALAN led to a 5% increase in growth rate, 67% in cell number and 49% in chlorophyll-a concentration, compared to the dark night condition. Exposure to red ALAN led to a similar response to the green ALAN, as it lead to higher chlorophyll-a, but it did not affect growth rate and total cell numbers compared to the dark treatment. Red and green ALAN treatments also affected the diatom-dominated phytoplankton assemblage. For example, red ALAN led to 118% and 80% higher total cell count and chlorophyll-a concentration compared to the dark control. More interestingly, red and green ALAN led to a similar assemblage response by balancing the biomass of the most abundant species (thus leading to higher evenness), but also by enhancing the biomass of the most abundant species relative to the subdominant species. These effects were less pronounced in response to the white ALAN treatment although it had a significant impact on assemblage richness.

Previous studies on the effect of white ALAN on freshwater primary producers have reported longer-term (6 week experiments) increases in the abundance of benthic microalgae (Hölker et al., 2015) but also shorter-term (3 weeks) decreases in periphyton abundance, as well as community composition shifts (Grubisic et al., 2017). Our experimental findings provide additional insights into ALAN effects by offering comparative information on different LED colours. Our findings show that although exposure to white ALAN can lead to a shift in diatom assemblage structure and biomass increase within 12 days of exposure, this effect was less pronounced compared to the red and green LED, and our white ALAN treatment had no effect on the growth rate of the green microalgae *Tetraselmis*.

Our findings also suggest that the red and green ALAN colours have the potential to enhance the growth of Harmful Algal Bloom (HAB) species such as the diatom *Skeletonema* sp.. This species is commonly known for forming dense blooms causing mortality to other organisms through physical damage (e.g. fish gill lesions) or anoxia, consequently impacting trophic interactions, biodiversity and overall ecosystem health (Kent, Whyte, & LaTrace, 1995). This finding is supported by Oh et al. (2008) who showed that green LEDs can selectively stimulate the growth of the diatom *Skeletonema costatum* compared to other species in the assemblage. The biomass enhancement under red and green ALAN of planktic colonial diatoms such as *Skeletonema* and *Thalasioseira* compared to epiphytic and epipsamic diatom genera might be attributed to an adaptation of these planktic forms to better exploit these colours which reach into deeper waters (Falcón et al., 2020) thus overcoming light limitation. Alternatively, this adaptation might have developed to enable planktic diatoms to sequester the dim moonlight which is richer in the warmer wavelengths (i.e. 4100K compared to 5000K of the sun) (Ciocca & Wang, 2013; Veilleux & Cummings, 2012).

A key question is why green and to some extent also red ALAN had a stronger effect on *Tetraselmis’s* growth compared to white light. A first insight stems from comparisons with previous studies that focused on maximizing the growth and biochemical composition of *Tetraselmis* to fully exploit the industrial potential of this algae. Unlike our 12:12 light period, these studies used continuous (24 h) high intensity LED illumination of different monochromatic LEDs (Abiusi et al., 2014; Aidar et al., 1994; Schulze et al., 2016). Abiusi et al. (2014) reported maximum growth and biomass concentration under red and white continuous light, whereas these traits were less pronounced under green light (all light conditions were standardised at 160 μmol m^−2^ s^−1^). Aidar et al. (1994) also reported increased growth under continuous red and white light compared to the blue-green light (all light conditions were standardised at 25 μmol m^−2^ s^−1^). These results contrast with the higher growth rate under green, dim night-time illumination found in our experiment. This discrepancy raises the question of whether dim light at night has the potential to induce different responses to LED wavelengths compared to higher intensity light at night.

An intriguing possibility is that photosynthetic organisms have developed higher sensitivity to night-time green-yellow light compared to UV, blue or far-red light. This could be explained by the fact that moonlight’s spectral peak is located at ~ 560 nm (Veilleux & Cummings, 2012), which is between green and yellow wavelengths. However, a green light receptor is yet to be identified in algae and flowering plants (Martin W Battle, Vegliani, & Jones, 2020). In fact, chlorophyll-a is widely recognised to show maximum absorbance in blue and red wavelengths (Fig. S1) (Clementson & Wojtasiewicz, 2019). Despite this, several recent studies have underlined the importance of other light wavelengths, including green, for microalgal biology (Ravelonandro et al.,2008; Mohsenpour et al,2012; Mattos et al., 2014; de Mooij et al. 2015; Gupta et al 2019; Ayalon et al. 2019) and plant growth (Kim, Goins, Wheeler, & Sager, 2004; Van, T K; Haller, W T; Bowes, G; Garrard, 1977). Although there is not known green light receptor in nature, other photoreceptor families that are involved in algae and plant responses to light may have played a role in our study. For instance, although cryptochromes are the primary receptors for UV-A and blue light, it has been reported that green light affects cryptochrome photochemistry and activity as green light reverts cryptochromes to their inactive state (Banerjee et al., 2007; Bouly et al., 2007). In particular, cryptochromes integrate green light signals into the circadian system as well as modulating plant growth and architecture in response to an increase in green/blue light ratio under a canopy (Martin W Battle et al., 2020; Martin William Battle & Jones, 2020; Trupkin, Karayekov, Buchovsky, Rossi, & Jose, 2020).

Another explanation could involve the phytochromes which are the red/far-red light receptors that control major developmental and growth events in land plants and algae including entrainment of the circadian clock and shade avoidance (Falcón et al., 2020; Rockwell et al., 2014). *Tetraselmis sp.* photocycle was shown to be controlled by the ratio of red/far-red (Rockwell et al., 2014). It is possible that in our study, exposure to natural white ALAN disrupted the circadian resonance of *Tetraselmis sp.,* whereas green (and to some extent red) LED light did not impair the circadian entrainment. This would have to be confirmed by monitoring chlorophyll abundance or clock regulated gene expression during a 24-hour period in presence of different ALAN conditions. While more mechanistic studies are needed in order to fully comprehend the responses of algae and plants to green light, our results contribute to the growing evidence that green light can profoundly affect their biology.

Our data show that there is a significant impact of green and red ALAN on phytoplankton that should be taken into account when planning nocturnal illumination in marine environments. In fact, these results may lead to conservation dilemmas, as both red and green LED lights have been suggested as alternative ALAN sources for public illumination. Specifically, red light illumination has been recommended in coastal areas because it interferes less with sea turtle nesting and hatching compared to broad spectrum white light (Miller and Bretschneider 2006). Similarly, the use of green light has been recommended to minimise the impact of light pollution on migratory birds (Poot et al. 2008). In general, shifting spectral signatures towards longer wavelengths than blue light seems to be beneficial to many organisms, including insects (Owens et al. 2020), bats (Spoelstra et al. 2017) and songbirds (Ouyang et al. 2017). However, our study shows that the use of green and red ALAN LEDs can impact aquatic primary producers by enhancing the growth of different taxonomic groups (green algae and diatoms) indicating a potential to encourage eutrophication phenomena in marine coastal (but potentially also freshwater) systems where these taxonomic groups are also present. Although the batch culture set-up used in our study is more representative of coastal systems affected by pulsed nutrient inputs (Spatharis et al. 2007), it would be interesting to also simulate systems that show less pronounced fluctuations using continuous or semi-continuous nutrient supply setups.

In conclusion, our study shows that ALAN can impact phytoplankton growth and assemblage structure, but also that this effect depends on the wavelength of light used, as green and red light accelerate growth and modify assemblage structure. While more experiments are needed to verify how generalizable these results are to other species, plankton communities and geographic areas, we stress that it is important to consider the colour of light in any future illumination of coastal areas.

## Supporting information

Diamantopoulou_et_al_Main.pdf

## Acknowledgements

This work was funded by “The A.G. Leventis Foundation” and “Nissad Development Company” (sponsors had no further involvement in the research). We further thank the IBAHCM Aquaria Staff for the technical support. We would also like to thank Sébastien Jubeau and Douglas McKenzie for their helpful suggestions in the initial stages of this study.

## Author contributions

CD, EC, DMD and SS designed the study. JC and NM designed the light system. CD, EC, DMD and SS performed the experiment. CD, EC and SS performed the lab analyses. SS and EC performed the statistical analyses. EK contributed to the interpretation of the results. CD, DMD and SS wrote the paper. All other authors read and commented on multiple drafts of the manuscripts, and all authors approved the final submitted version.

## Data availability

We will upload our data to the Dryad repository should this manuscript become accepted.

