## Supplementary material for "Wavelength-dependent effects of artificial light at night on phytoplankton growth and community structure": Diamantopoulou_et_al_Main.pdf

### Supplementary tables

**Table S1**. Pairwise comparison of growth rate, and cell number (cells/ml), and chlorophyll a concentration (µg/L) between the ALAN treatments, showing the effect size between treatment levels and corresponding p-values. Note that cell number and chlorophyll a were measured at day 18 of the experiment.

| Pairwise comparison | Growth rate | Cells | Chl a |
|---|---|---|---|
| **Dark - Green** | -0.142 (0.005) | -65867 (0.002) | -17.55 (<0.000) |
| **Dark - Red** | -0.080 (0.086) | -28200 (0.149) | -9.01 (0.010) |
| **Dark - White** | -0.035 (0.427) | 17933 (0.350) | 4.83 (0.137) |
| **Green - Red** | 0.062 (0.178) | 37667 (0.060) | 8.54 (0.014) |
| **Green - White** | 0.178 (0.001) | 83800 (0.000) | 22.38 (<0.000) |
| **Red - White** | 0.074 (0.017) | 46133 (0.025) | 13.85 (<0.001) |

**Table S2.** Pairwise comparison of diatom assemblage total cells, Menhinick richness, evenness, chlorophyll a and c between the ALAN treatments, showing the effect size between treatment levels and corresponding p-values. Note that all estimates are based on data from day 12 of the experiment (see methods section for details).

| Pairwise comparison | Total cell count | Menhinick | Evenness | Chl a | Chl c |
|---|---|---|---|---|---|
| **Dark - Green** | -1003 (0.004) | 0.068 (0.048) | -0.089 (0.001) | -3.73 (0.004) | -0.545 (0.230) |
| **Dark - Red** | -1560 (<0.000) | 0.083 (0.018) | -0.129 (<0.000) | -6.61 (<0.000) | -1.425 (0.005) |
| **Dark - White** | -849 (0.012) | 0.086 (0.015) | -0.069 (0.008) | -4.02 (0.002) | -0.878 (0.064) |
| **Green - Red** | -557 (0.077) | 0.0157 (0.627) | -0.041 (0.097) | -2.88 (0.018) | -0.881 (0.063) |
| **Green - White** | 154 (0.609) | 0.01846 (0.569) | 0.019 (0.413) | -0.29 (0.794) | -0.334 (0.461) |
| **Red - White** | 710 (0.028) | 0.003 (0.934) | 0.060 (0.081) | 2.59 (0.030) | 0.547 (0.233) |

### Supplementary figures

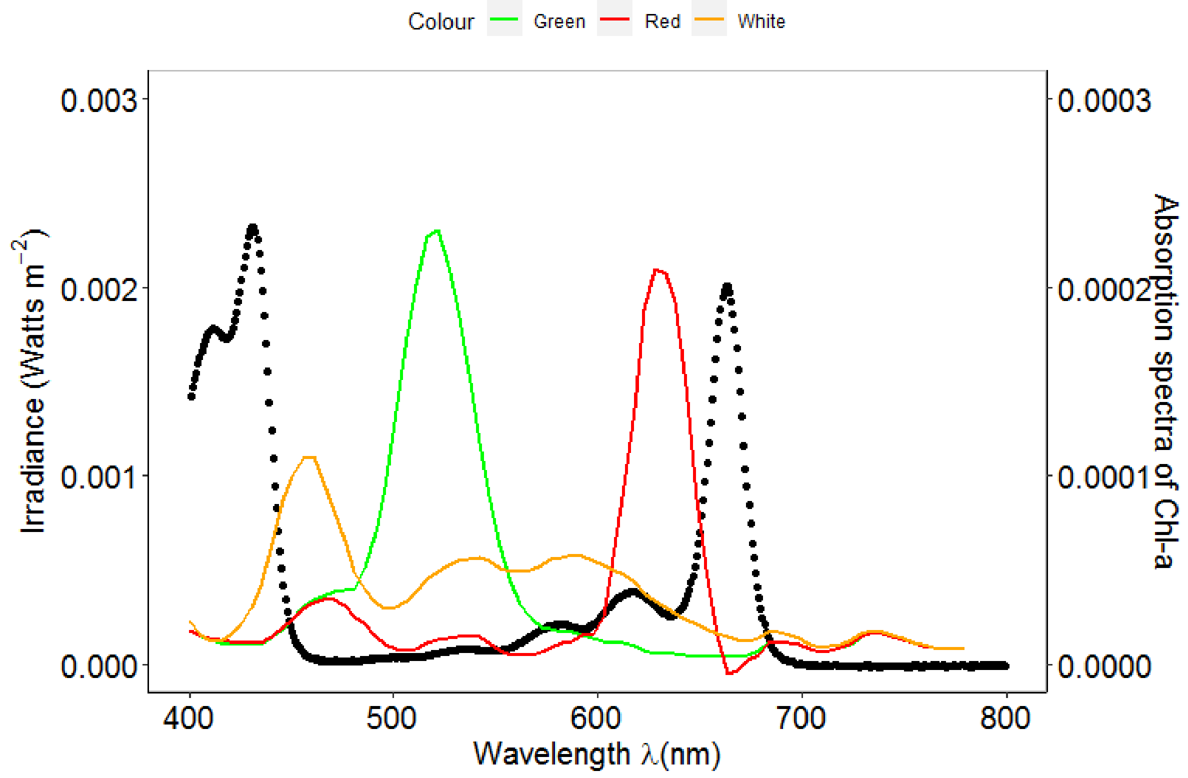


**Figure S1.** Spectral profiles of the ALAN treatments (solid coloured lines) and absorption spectra of Chlorophyll a (black dotted line) (Clementson and Wojtasiewicz 2019). The irradiance level of all ALAN conditions was standardised at 0.023 Watt $m^{-2}$ in all ALAN treatments.

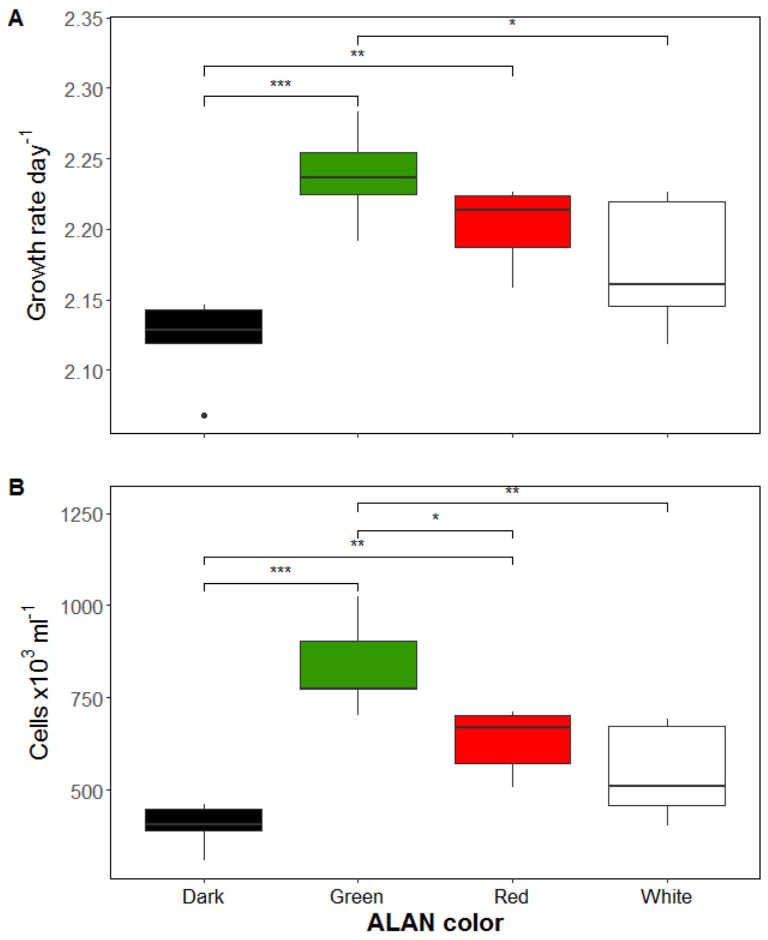


**Figure S2.** Effect of ALAN treatments (Dark, Green, Red and White) on the growth rate calculated during the exponential growth phase (panel A) and on the cell concentration (panel B). The data was obtained from cell counts from a previous pilot experiment to the experiment discussed in the main text of this paper. Here, all the ALAN sources were standardised at an intensity of 20 lux (this corresponded to an irradiance of 0.080 Watt $m^{-2}$ for the red ALAN, 0.016 Watt $m^{-2}$ for the green ALAN, and 0.029 Watt $m^{-2}$ for the white ALAN). The light colours were allocated in a different randomised order in the experimental boxes compared to the main experiment. We are showing these data here to clarify that: i) the results of the two experiments are comparable irrespective of the type of light standardization used (irradiance in Watt $m^2$ or illuminance in Lux); ii) the results are also not affected by the box used for each LED colour. Pairwise comparisons show differences between treatments (not shown: $p > 0.05$, *: $p <= 0.05$, **: $p <= 0.01$, ***: $p <= 0.001$).
